# Genetic diversity analysis of Chinese plum (*Prunus salicina* L.) based on whole-genome resequencing

**DOI:** 10.1101/2020.08.28.271411

**Authors:** Xiao Wei, Fei Shen, Qiuping Zhang, Ning Liu, Yuping Zhang, Ming Xu, Shuo Liu, Yujun Zhang, Xiaoxue Ma, Weisheng Liu

## Abstract

Chinese plum (*Prunus salicina* L.), also known as Japanese plum, is gaining importance due to their extensive genetic diversity and nutritional attributes beneficial for human health. Single-nucleotide polymorphisms (SNPs) are the most abundant form of genomic polymorphisms and are widely used in population genetics research. Here, we construed high-density haplotype map by whole-genome resequencing of 67 *Prunus* accessions with a depth of ~20× to evaluate the genome-level diversity and population structure. The phylogenetic analysis, the principal component analysis, and the population structure profiling, indicated that the 67 plum accessions could be classified into four groups corresponding to their origin location, the southern cultivar group (SCG), the northern cultivar group (NCG), the foreign cultivar group (FG), and the mixed cultivar group (MG). Some cultivars from South China were clustered with the other three groups. The genetic diversity indices including the private allele number, the observed heterozygosity, the expected heterozygosity, and the nucleotide diversity of the SCG were higher than those of the NCG. The gene flow from the SCG to the FG was detected as well. We concluded that the origin center of Chinese plum was at the Yangtze River Basin in South China. This study provided genetic variation features and population structure of Chinese plum cultivars, laying a foundation for breeders to use diverse germplasm and allelic variants for improving Chinese plum varieties.

## 1. Introduction

Plum is one of the most important fruit crops in the world, possessing extensive genetic diversity and high economic value. The largest plum producer is China, with an annual production of 6,801,187 metric tons in 2018, accounting for 53.9% of the world’s total [1]. Within the genus, *Prunus*, Chinese plum *(Prunus salicina* L., 2n = 2x = 16), also known as Japanese plum, is widely grown for fresh market consumption and the canning industry, which includes both pure Chinese plum and its hybrids with other diploid plum species, such as *Prunus simonii* Carr., *Prunus cerasifera* Ehrh., *Prunus americana* Marsh., and others [2].

Chinese plum possible originates from the Yangtze River Basin, and now there are abundant plum germplasm resources in China [3]. With its long growing history and extensive geographical distribution, there are more than 800 indigenous plum cultivars in China derived from *Prunus salicina* L. alone. Over 600 of them are now preserved at National Germplasm Repository for Plums and Apricots (NGRPA) located in Xiongyue, Liaoning Province, China [4,5]. Yu et al. [6] carried out comprehensive phenotyping of 405 Chinese plum cultivars and its hybrids from NGRPA, totally 32 morphological and agronomic characters were investigated, of which their coefficients of variations (CVs) were calculated ranging from 14.85% to 47.09%, suggesting that the genetic variability of Chinese plums distributed widely.

Genetic variability is the prerequisite for any plant breeding program. Learning the extent and structure of genetic variation in germplasm collections is a crucial step for the efficient conservation and utilization of biodiversity in cultivated crops, and using diverse plum resources to broaden the genetic base of worldwide plum cultivars is the critical objective for plum breeders [2,7]. Previous efforts have been made to understand the genetic diversity of plum better. Zhang and Zhang [8] took the lead in collecting plum germplasm resources and evaluating their genetic diversity on the basis of morphological traits and isozyme polymorphisms, whereas, morphological traits were highly susceptible to environmental factors, so the estimates of genetic diversity were not precise; while isozymes had a low degree of polymorphism and hence were not efficient enough for the characterization of germplasm genetic diversity [9]. Overcoming the above-mentioned problems, DNA-based markers became widely applied in plum genetic diversity analysis, including random amplified polymorphic DNAs (RAPDs) [5,10], simple sequence repeats (SSRs) [11–13], inter-simple sequence repeats (ISSRs) [2,14]. In addition, benefiting from the advances in next-generation sequencing (NGS) technologies, genotyping by sequencing (GBS), provides a great wealth of information that makes it possible to identify thousands of single nucleotide polymorphisms (SNPs), which, after adequate filtering, allow us to carry out detailed genetic diversity studies [7,15–17].

By conducting DNA-based marker assessments, Chinese plums were generally classified into two major groups, the southern cultivar group (SCG) and the northern cultivar group (NCG) [18–20]. The *P. salicina* system in South China was rather primitive than that in North China basing on a palynology study, and the spread of *P. salicina* was from the south to the north [21]. However, simply cultivar clustering works are inadequate for the integrated study of genetic diversity of Chinese plum. As described in peach, large scale SNP data-based diversity analysis boosted the deciphering of the evolution and domestication of peach germplasm resources [22,23].

In the present study, we used SNP markers generated from high depth whole-genome resequencing data (an average depth of ~20×) for the first time to elucidate the pattern of genetic diversity, population structure, and domestication of a diverse *P. salicina* collection. An in-depth understanding of their genetic relationships could benefit plum germplasm conservation and utilization, and lead to new cultivar improvement.

## 2. Results

### 2.1 Markers development based on whole-genome resequencing

We collected 67 plum accessions representing different geographic and morphological characteristics (Table S1). The 67 plum accessions were sequenced using the Illumina HiSeq 2500 platform with a sequence depth higher than 20×; a totally 462.6 Gb clean data were retained with an average of 6.9 Gb for each accession after filtering out low-quality reads. The quality of the sequencing data was high with a Q30 > 85%, and a GC content slightly fluctuated around 38.0%. The mapping rate to the peach genome was range from 85.73% to 88.57%, with a average of 87.49% (Table S1). We called SNPs with the unique mapped reads, and a total of 16,600,033 SNPs and 2,107’664 INDELs were identified across the 67 accessions using SAMtools v1.4 software. After filtering out these in low quality, we obtained 14,549,234 SNPs which could ensure the accuracy and reliability of subsequent genetic diversity and population structure analyses (Figure 1). About 43.81% of the SNPs were located in intergenic regions, and 11.88% were in coding regions. The non-synonymous to synonymous substitution ratio (dN/dS) for the SNPs in the coding regions was 15.92. Transitions were found in a proportion of 59.58% (10315764/16600033), with a transition/transversion ratio (Ts/Tv ratio) of 1.47 (Figure 1, Table 1).

**Figure 1.**
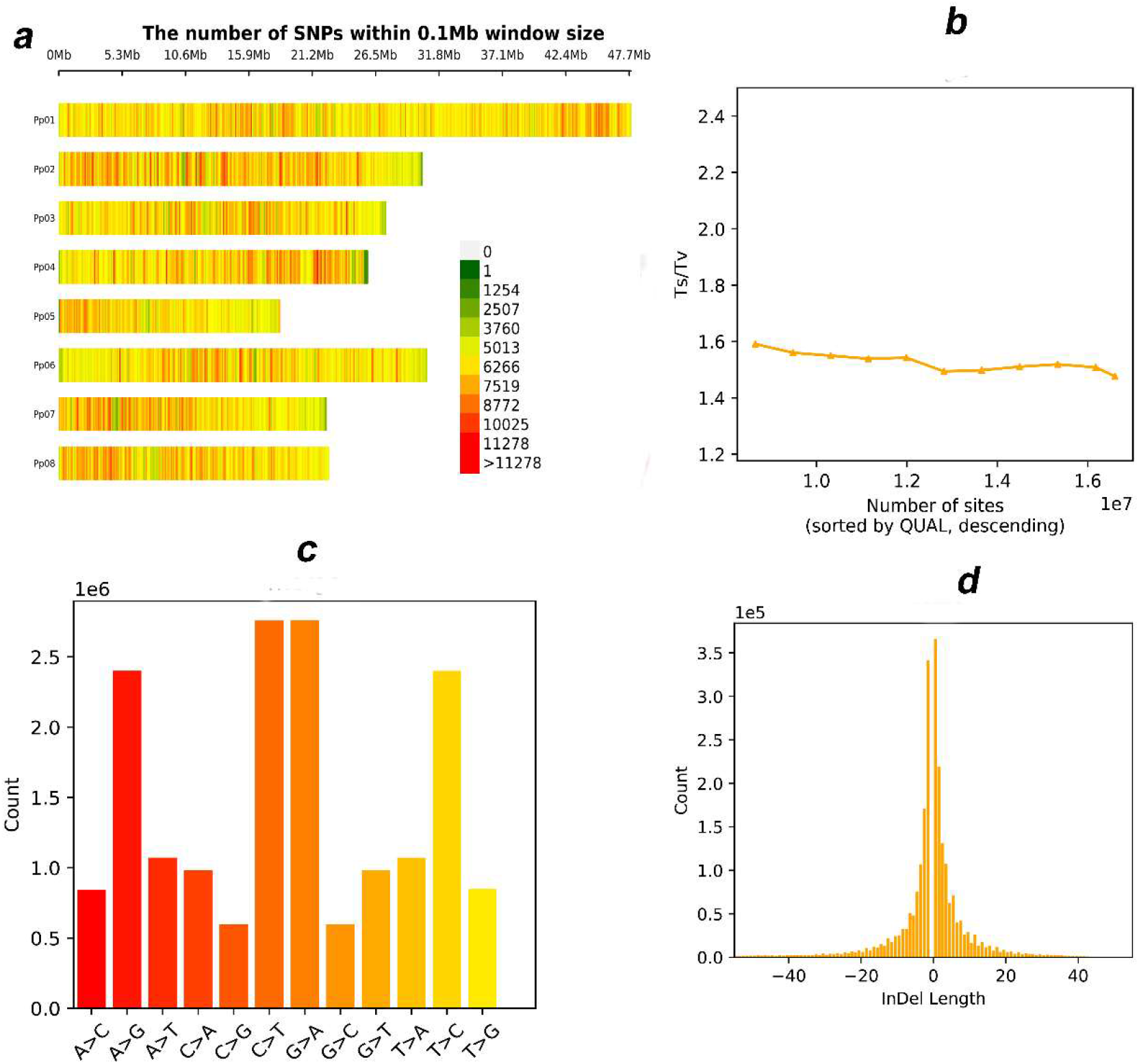
The statistics of markers generated from the whole-genome resequencing of the 67 plum accessions. a) The chromosome-scale SNP distribution; b) The distribution of transition/transversion ratio (Ts/Tv ratio); c) The counts of different types of transitions and transversions; d) The counts of genome-wide insertions and deletions.

**Table 1.** Statistics of variations across different chromosomes

| Categories | Numbers | Ratio |
| --- | --- | --- |
| Raw SNPs | 16,600,033 | 88.73% |
| InDels | 2,107,664 | 11.27% |
| Filtered SNPs | 14,549,234 | 87.65% |
| Intergenic regions SNPs | 7,272,007 | 43.81% |
| Coding regions SNPs | 1,971,748 | 11.88% |
| Transition | 10,315,764 | 59.58% |
| Transversion | 6,999,126 | 40.42% |
| Transition/Transversion | -- | 147.00% |
| Non-synonymous SNPs | 1,855,201 | 94.09% |
| Synonymous SNPs | 116,547 | 5.91% |

### 2.2 Phylogenetic analysis

To exploit the relationship among the Chinese plum, a Neighboring-joining phylogenetic tree of the 67 accessions was constructed with all SNPs (Figure 2a). The phylogenetic tree classified the accessions into four main groups which are corresponding to their origin location: (1) the southern cultivar group (SCG) was comprised of the plum cultivars from Sichuan, Guizhou, Yunnan, Guangdong, Guangxi, Zhejiang, and Fujian Province; (2) the northern cultivar group (NCG) comprised of plum cultivars mainly from Hebei, Henan, Shandong, Shaanxi Province, and the south part of Liaoning Province; (3) the foreign cultivar group (FG) included plum cultivars from the USA and Japan; and (4) the mixed cultivar group (MG) consisted of newly bred cultivars from the NGRPA and several cultivars originated SCG and FG. Notably, we fould ‘Saozouli’ and ‘Shuili’ cultivars collected from south China were clustered with NCG, ‘Zaohuangli’ and ‘Jinshali’ with FG, and ‘Wanshu huanai’, ‘Huahongli’ and ‘Abazhoumeiguili’ with MG, suggesting that SCG is more diverse than any other group.

**Figure 2.**
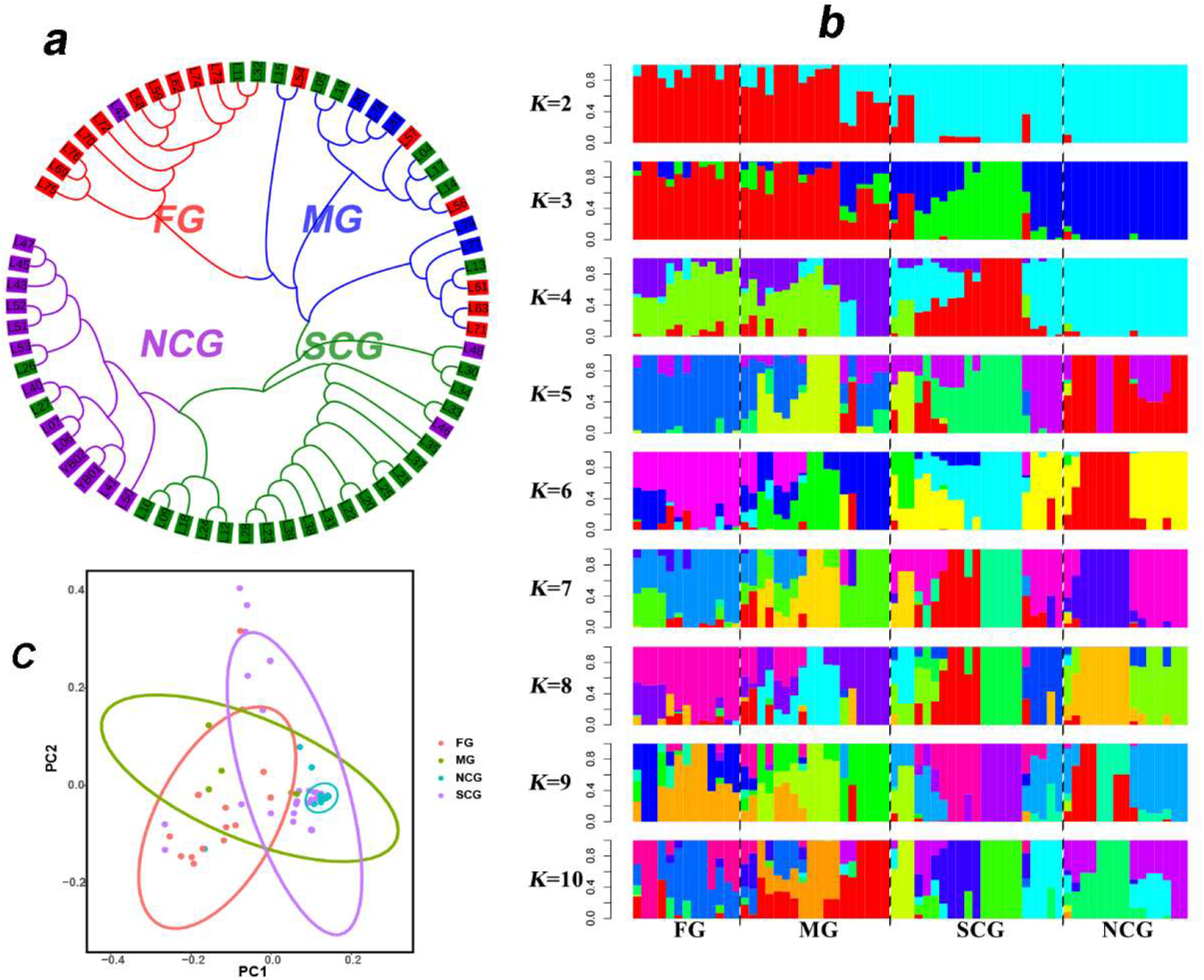
The population structure of the 67 plum accessions. a) Neighboring-joining phylogenetic tree constructed using SNPs at fourfold degenerate site. Each group was color coded; b) Bayesian model-based clustering of the 67 plum accessions with the number of ancestry kinship (K) from 2 to 10; c) Principle Component Analysis (PCA) of the 67 plum accessions. FG, the foreign cultivar group; MG, the mixed cultivar group; SCG, the southern cultivar group; NCG, the northern cultivar group.

### 2.3 Population structure analysis

Population genetic structure analysis was performed based on high-quality SNPs. We employed 5-fold cross validation to inference the number of ancestor population, K (Figure 2b). The NCG was divergent from SCG when the K >2, suggesting the NCG was derived from SCG. We observed that SCG includes individuals with similar or different genetic structure to these from other groups like NCG and FG, further indicating that it has higher diversity than others. The observation was further supported by genetic diversity analysis with π (Figure 3b). Thus, other groups including FG and NCG potentially associated with SCG during plum evolution.

**Figure 3.**
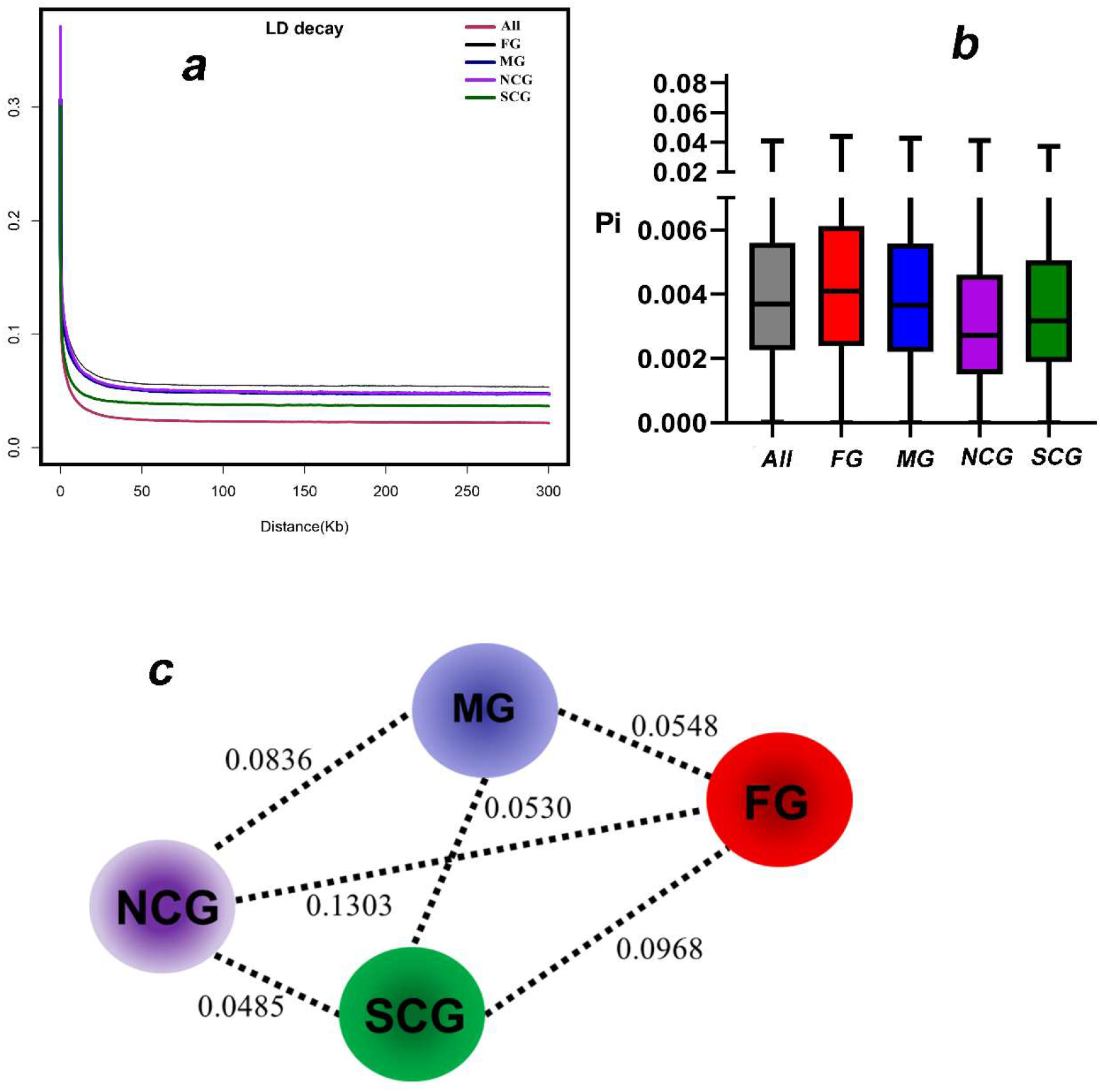
The genetic diversity of different groups. a) The decay of linkage disequilibrium (LD) measured as the squared correlation coefficient (*r*^2^) by pairwise physical distance; b) The nucleotide diversity (π) of different groups; c) The genetic differentiation analysis between groups. The values between pairs indicate population divergence (*F*_ST_). FG, the foreign cultivar group; MG, the mixed cultivar group; SCG, the southern cultivar group; NCG, the northern cultivar group.

### 2.4 Principal component analysis

To further confirm the relationship among the cultivars, we performed principal component analysis (PCA) toward the 67 plum accessions (Figure 2c). Based on the principal component plot with first two principal components, the NCG exhibited a relatively close relationship with SCG, which was consistent with the phylogenetic tree and population structure analysis. We deduced that the NCG could be considered as a more independent subgroup of SCG and may be derived from a specific ecotype. Furthermore, samples from SCG showed some degree of correlation, suggesting the close relationship between SCG and others. The results further confirm that other groups potentially originate from SCG.

### 2.5 Genetic diversity analysis

As shown in Table 2, the expected heterozygosity (He) of the *Prunus* populations varied between 0.234 and 0.304; the observed heterozygosity (Ho) of the *Prunus* populations ranged between 0.328 and 0.429; Wright’s F-statistic (*F*_IS_) of the *Prunus* populations varied between −0.241 and −0.146; the number of private alleles (A_P_) in the *Prunus* populations varied between 546 and 31,285; the nucleotide diversity (π) ranged between 0.00358 and 0.00467. The mean nucleotide variation of *P. salicina* was higher than that of other perennial crops, such as peach (π = 0.0015) [24], cassava (π = 0.0026) [25], and apricot (π = 0.0027) [26], but lower than that of date palm (π = 0.0092) [27]. The Tajima’s D values of the four groups were all tested positive (1.002 – 1.497) and significantly different from zero. Thus, the null hypothesis of neutral evolution could be rejected. As shown in Figure 3a, the LD decay rate was fastest in the SCG and was lowest in the FG, suggesting a lower frequency of genetic recombination in the FG, as they included artificially hybridized cultivars.

**Table 2.** The statistical values of genetic diversity within different populations.

| Groups | Private allele number (Ap) | Observed heterozygosity (Ho) | Expected heterozygosity (He) | Nucleotide diversity ($\pi$) | Wright's F-statistic ($F_{IS}$) | Tajima's D |
| --- | --- | --- | --- | --- | --- | --- |
| FG | 31,285 | 0.4290 | 0.3036 | 0.00467 | -0.2411 | 1.4974 |
| MG | 30,112 | 0.3658 | 0.2846 | 0.00433 | -0.1560 | 1.0094 |
| SCG | 27,287 | 0.3365 | 0.2606 | 0.00395 | -0.1460 | 1.0022 |
| NCG | 546 | 0.3280 | 0.2336 | 0.00358 | -0.1829 | 1.3211 |

### 2.6 Genetic differentiation analysis

The *F*_ST_ between the four groups varied between 0.0485 and 0.1303 (Fig. 3). The *F*_ST_ value of MG-FG, MG-SCG, and SCG-NCG pairs was relatively lower among all the group-pairs analyzed, indicating that the genetic differences within populations were higher than those between populations and the possible genetic exchange between populations. The *F*_ST_ value between the NCG and FG groups were the largest among all the groups analyzed, which may have been due to geographical isolation, low gene flow between populations, and significant genetic differences.

### 2.7 Gene flow analysis and inferred evolutionary path

We analyzed the gene flow between the four geographic groups (Figure 4b, 4c). Allowing one or two migration events (m = 1 or 2), we observed that gene flow occurred between the SCG and FG accessions, which likely reflected their many shared genomic components due to hybridization in their domestication and breeding histories. Considering the geographical locations of SCG and NCG, though these two ecological groups were close to each other, we found no gene flow between them, indicating a relatively independent domestication process.

**Figure 4.**
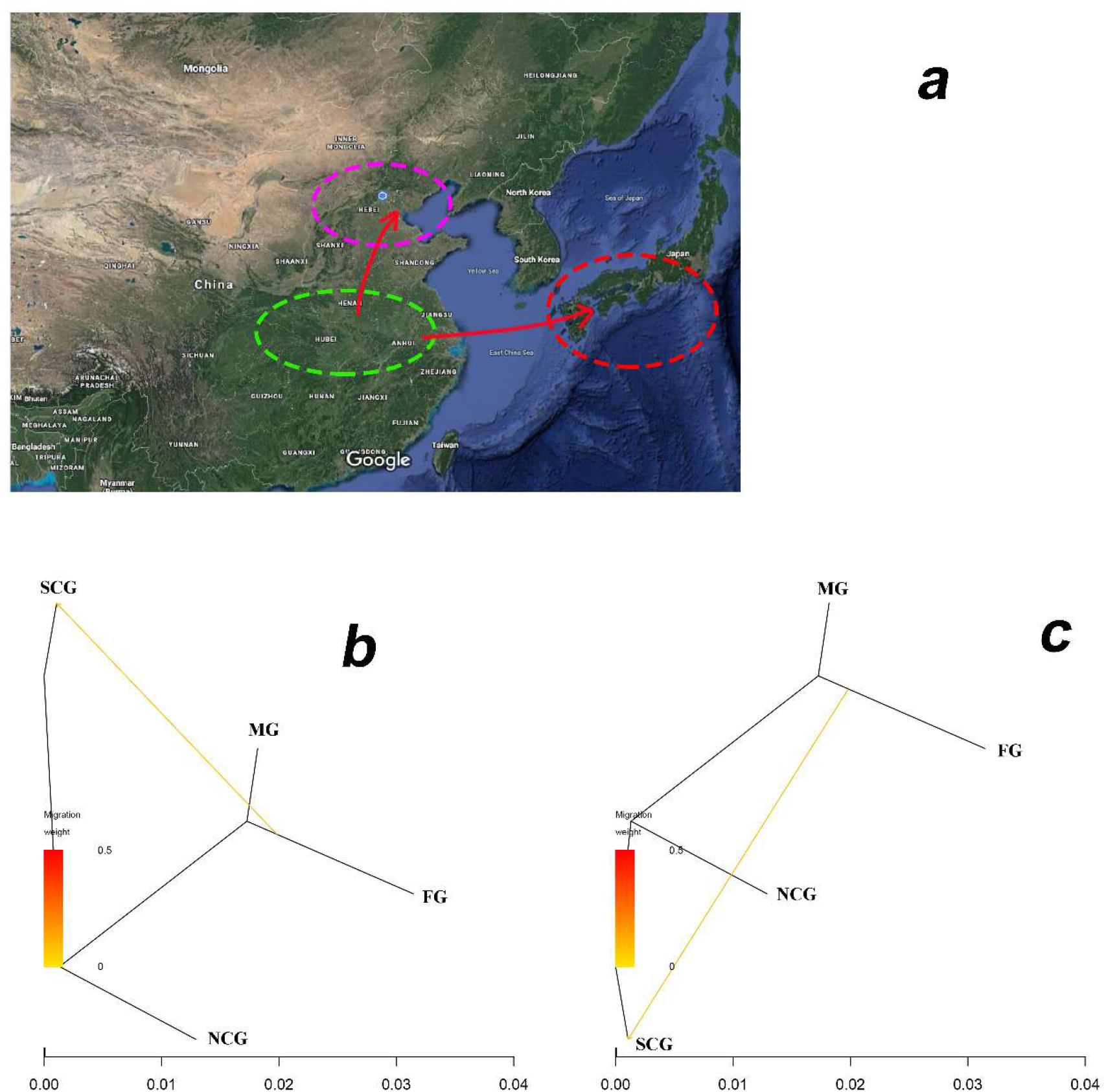
The inferred evolutionary path (a) and gene flow analysis between populations as inferred by TreeMix using a model with one (b) and two (c) admixture events. Admixtures are colored according to their weight. FG, the foreign cultivar group; MG, the mixed cultivar group; SCG, the southern cultivar group; NCG, the northern cultivar group.

## 3. Discussion

SNPs have become tremendously important as markers for genetics research in plants because they have been found to be in high frequency, display a lower mutation rate compared to SSR-based markers, and they are uniformly distributed across the genome [28]. NGS is a kind of low-cost technologies to discover large numbers of SNPs for extensive genetic studies, including within-species diversity analysis, linkage map construction, and genome-wide association studies (GWAS), which is allowing significant advances in plant genetics and breeding [6,22,23,29]. The genus *Prunus* has shown conserved intraspecific and intragenic collinearity in the *Rosaceae* family, and peach has been considered as a model species for the genus *Prunus* regarding multiple genetic types of research [30–33]. On these bases, for the first time, we conducted high-depth (~20×) whole-genome resequencing of 67 plum accessions using the peach genome sequence as a reference, obtaining an average of 6.9 Gb high-quality data for each acquisition, which ensured the accurate and comprehensive population genetic analysis.

Based on the high-quality SNPs, the 67 plum accessions in this study were divided into four groups, the SCG, NCG, FG, and MG, supported by the phylogenetic tree, the population structure analysis, and the PCA results, which were partly in accordance with the previously obtained results through RAPD, ISSR, and SSR marker assessing [18–20]. We noticed that several cultivars from the south part of China distributed in other three groups (Figure 2a), combining with the fact that the *P. salicina* system in South China was rather primitive than that in North China basing on a palynology study, we speculated that, for SCG, the genetic background was relatively broader, the gene exchange was more frequent with other groups, and the possible origin center of Chinese plum was at the Yangtze River Basin in South China [3,20,21,34]. For the five newly bred cultivars from the NGRPA in MG, one of their parents was from the USA (‘Blackamber’, ‘Friar’) or Japan (‘Akihime’) and the other parent was native to China, and the gene flow between SCG and FG was detected (Figure 4). Thus, we assumed that those cultivars in MG were genetically related. As mentioned in other studies, most of the Chinese plum cultivars from Japan and the improved Chinese plum hybrids from the USA were found distributed across Chinese indigenous plum group through dendrogram analysis [2,5,20], which was according with the *P. salicina*-predominant genetic components of improved Chinese plum hybrids [35], and in agreement with the fact that China-originated plum cultivars were initially introduced to Japan, and then imported to the USA in 1870 from Japan by Luther Burbank [3].

Southwest China was speculated as a primitive domestication center of plum as there were wild plum populations discovered in Yunnan, Sichuan, and Guizhou Province [20]. Suggesting by the high Shannon index of diversity, the high value of average sufficient allele, and the high expected heterozygosity, the Chinese plums cultivated in southwest China were tested with high genetic diversity using SSR markers [20]. Additionally, by analyzing the phenotyping variation of 405 plum accessions distributed in different geographical regions in China, the fruit weight of cultivars in South China exhibited the most significant coefficient of variation [6]. The π, Ho, and He of the SCG calculated with SNPs in our study were higher than those of the NCG, corresponding with the higher genetic diversity of the SCG; these genetic diversity indices of the SCG were lower than that of the FG and MG, presumably due to the artificial hybridized cultivars in the FG and MG.

Generally, the genetic differentiation between populations is considered moderate when the *F*_ST_ value is higher than 0.05 and is considered highly differentiated when the *F*_ST_ value is higher than 0.15 [36]. In our study, the NCG-FG pair possessed the highest *F*_ST_ value, following by the SCG-FG couple. In contrast, the *F*_ST_ value of the MG-FG was significantly lower as there were five newly selected plum cultivars by NGRPA in the MG with foreign lineage from the FG. Human intervention via the artificial selection of favorable phenotypic traits to enhance production and improve desirable agronomic traits can both reduce the levels of genetic variability and skew allele frequencies [37].

## 4. Materials and Methods

### 4.1 Plant materials

A diverse collection consisting of 67 *Prunus* spp. accessions, including 65 *P. salicina* accessions and two *P. simonii* accessions, were selected for the whole-genome resequencing study (Table S1). All these trees were held at NGRPA located in Xiongyue county, Liaoning Province, China (40°18’ N, 122°16’ E) in a planting density of 3.0 × 4.0 m, trained to the open vase system with three or four main branches, and maintained under conventional management and pest control operations.

### 4.2 DNA extraction, library preparation, and sequencing

Genomic DNAs were isolated from the young and healthy leaf samples of the 67 accessions using a modified CTAB protocol [38]. The quality and integrity of the DNAs were examined using a NanoDrop^®^ spectrophotometer (ND-1000, Thermo Fisher Scientific Inc., USA) followed by electrophoresis on 1% agarose gels. Quantification of the DNA samples was assessed with Qubit™ (2.0 Fluorometer, Invitrogen, Carlsbad, CA, USA).

High molecular weight DNA aliquots with 230/260 and 260/280 ratios ranging between 1.8 – 2.0 and 1.8 – 2.2, respectively, were then sent to BGI (Shenzhen, China) for library construction and sequencing. The insert-size of the libraries was 500 bp, and the length of pair-end reads was 150 bp. All the libraries were sequenced using the Illumina HiSeq 2500 platform (Illumina, San Diego, CA, USA).

### 4.3 Reads mapping, SNP calling, and SNP annotation

The qualified paired-end reads of each accession were aligned against the peach reference genome v2.0 [24] using BWA v0.7.12-r1039 with the parameters mem -t 4 -k 32 -M [39], and SNPs were identified using SAMtools v1.4 [40]. Low-quality SNPs were filtered out by minimum minor allele frequency (mnMAF < 0.01) and missing data per site (MDpS > 10%), and finally converted into Variant Call Format file (VCF). Gene-based SNP annotation was performed using the ANNOVAR package v2018-04-16 [41]. Based on the reference genome annotation, SNPs were categorized as occurring in exonic regions (overlapping with a coding exon), intronic regions (overlapping with an intron), upstream and downstream regions (within a 1 kb region upstream or downstream from the transcription start site), or intergenic regions. Additionally, the heterozygosity was calculated using VCFtools v0.1.14 [42].

### 4.4 Population structure analysis

TreeBeST v1.9.2 software was used to calculate the distance matrix [43]. RAxML v8.2.12 was used to construct the ML phylogenetic tree, and 1000 bootstrap replicates were used [44]. The obtained phylogenetic tree was visualized using MEGA v5.05 [45].

The principal component analysis was performed using the PLINK v1.07 software with default parameters [46].

We used ADMIXTURE v1.23 to infer the population structure [47]. To identify the best genetic clusters *K*, cross-validation error was tested for each *K* value from 2 to 10. The termination criterion was 1e-6 (stopping when the log-likelihood increased by less than 1e-6 between iterations).

### 4.5 Linkage disequilibrium (LD) analysis

Linkage disequilibrium (LD) was calculated using SNPs with MAF greater than 0.05 through PLINK v1.07 software with the following setting: --file --r2 --ld-window 99999 --ld-window-kb 200 --out. LD decay was calculated based on the squared correlation coefficient (*r*^2^) values between two SNPs and the physical distance between the two SNPs [46].

### 4.6 Gene flow analysis

The TreeMix v1.13 software was used to evaluate the gene flow among different groups with the parameters -se-bootstrap-k 1000 -m, where the number (-m) varied from one to three [48].

### 4.7 Genetic diversity and differentiation analysis

To evaluate the genetic diversity and differentiation, we used a 100-kb sliding window with a step size of 10 kb to calculated the number of private alleles (A_P_), the observed heterozygosity (Ho), the expected heterozygosity (He), the nucleotide diversity (π), the Tajima’s D value, Wright’s F-statistic (*F*_IS_), and the population-differentiation statistics (Fixation index, *F*_ST_) using VCFtools v0.1.14 [42].

## 5. Conclusions

In the current study, using high-quality SNPs generated from high-depth whole-genome resequencing of 67 *Prunus* accessions, we concluded that the origin center of Chinese plum was at the Yangtze River Basin in South China. This study also provided genetic variation features of plum cultivars and dissected their genetic diversity and population structure, laying a foundation for breeders to use diverse germplasm and allelic variants towards developing improved Chinese plum varieties.

## Data availability

All of the raw reads of the plum accessions generated in this study have been deposited in the public database of National Center of Biotechnology Information under PRJNA659814.

## Supplementary Materials

Supplementary material can be found at http://www.

## Authors’ contributions

ZQP and LWS conceived and designed the experiments. WX and SF performed the experiments. WX, SF and ZQP analyzed the data and wrote the paper. LN, ZYP, XM, LS, MXX and ZYJ collected samples and managed the materials. LWS revised the manuscript. All authors have read and approved the final manuscript.

